# Constraints and adaptations in crocodyliform skull evolution

**DOI:** 10.1101/2025.07.09.663933

**Authors:** Ananth Srinivas, Jen A. Bright, John A. Cunningham, Sandra A. S. Tavares, Fresia Ricardi Branco, Ismar de Souza Carvalho, Fabiano Vidoi Iori, Emily J. Rayfield

## Abstract

Crocodyliformes display a diverse range of skull morphologies though their evolutionary history. Extant crocodilians possess platyrostral (broad and flat) snouts, thought to be sub-optimal for resisting feeding loads due to the conflicting demands of feeding and hydrodynamic constraints. In contrast, numerous Mesozoic crocodyliformes possessed oreinirostral (dome-shaped) skulls, were terrestrial and hence free from hydrodynamic constraint. This study aims to examine the role of function in determining skull shape in crocodyliformes, and assesses the difference in stress resistance between oreinirostral and platyrostral taxa. We hypothesise that in the absence of hydrodynamic constraints, oreinirostral taxa have skulls that are better suited for resisting feeding induced loads. Using finite element analysis (FEA), we evaluated biomechanical performance in oreinirostral notosuchian taxa *Baurusuchus salgadoensis*, *Montealtosuchus arrudacamposi* and *Caipirasuchus paulistanus*; compared to the extant platyrostral *Alligator mississippiensis, Crocodylus niloticus,* and *Paleosuchus palpebrosus*. Results show that oreinirostral morphologies are comparatively better suited for resisting forces generated during feeding, with increased muscular efficiency, supporting the hypothesis that hydrodynamic constraints influence crocodyliform skull evolution.

## Introduction

Crocodilians, one of only two living groups of archosaurs, are represented by 28 extant species comprising of alligators, caimans, crocodiles, and gharials. They are considered conservative in their ecological and cranial disparity **(Gignac et al., 2019, Stubbs et al., 2021)**. Modern crocodilians are large semi-aquatic apex predators capable of high bite forces, related to numerous skull modifications such as closure of the antorbital fenestra, a relatively small braincase, and the formation of the secondary palate, which confers additional strength to the skull **(Preuschoft and Witzel, 2002, Rayfield and Milner, 2008)**. Numerous studies on crocodilian skull evolution have highlighted significant variation in snout proportions **(Busbey, 1995, Brochu, 2001, Pierce et al., 2008, 2009)**, offering valuable insights into ecological and biomechanical adaptations. This includes the repeated convergent evolution of longirostrine morphologies, which reflect specialised feeding strategies and habitat use **(Walmsley et al., 2013, McCurry et al., 2017a, McCurry et al., 2017b, Ballell et al., 2019)**. Understanding these patterns provides a framework for exploring the evolutionary pressures that have shaped crocodyliform diversity over time.

Crocodilian skulls are generally classified based on snout length, as longirostrine (long-snouted) and brevirostrine (short-snouted), but can also be classified as platyrostral (flat and dorsoventrally compressed) and oreinirostral (dome shaped and mediolaterally compressed) **(Busbey, 1995)**. Busbey (1995) evaluated the functional and mechanical implications of skull shape evolution in crocodilians using beam theory to link shape differences to diet and feeding strategies. Oreinirostral skulls, which are taller than they are wide, are considered better suited to resisting dorsoventral mechanical loads than platyrostral skulls. Skull flattening is thought to lead to a sub-optimal condition for resisting feeding loads, and likely evolved in extant crocodilians to reduce the effects of drag in the aquatic environment, or to gain higher resistance during torsional loads generated during feeding **(Busbey, 1995)**. Thus, extant crocodilians face competing functional demands between feeding efficiency and hydrodynamic performance **(McHenry et al., 2006, Rawson et al., 2024)**.

In contrast to extant crocodilians, crocodylomorphs exhibit a broader range of cranial morphotypes. During the Mesozoic, crocodyliformes occupied various ecological niches, with several protosuchians and notosuchians radiating onto land; and thalattosuchians, pholidosaurs and dyrosaurids occupying the oceans **(Pol, 2005, Sereno and Larsson, 2009, Stubbs et al., 2021)**. Notosuchians in particular were an extremely diverse group of terrestrial crocodyliformes spanning across the Jurassic and Cretaceous deposits of Gondwana, with certain sebecosuchians surviving till the Middle Miocene, about 11 million years ago. They exhibited several distinctive characteristics such as erect posture **(Pol, 2005)**, heterodont dentition **(Marinho and Carvalho, 2009)**, complex jaw movements to process food **(Ősi, 2014)**, and a wide range of diets including hypercarnivory, omnivory and herbivory **(Melstrom and Irmis, 2019, Borsoni et al., 2024)**. Numerous terrestrial notosuchians possessed oreinirostral skulls with feeding strategies such as ‘biting and slashing’, differing from yawing and the classic ‘death roll’ mechanism exhibited by their extant counterparts, performed to dismember large prey **(McIlhenny, 1935, Pooley and Gans, 1976, Fish et al., 2007)**. Herbivorous notosuchians had taller skulls, shorter snouts, and likely produced relatively higher bite forces for processing tough plant material, with similar adaptations found in certain lizards **(Aerts et al., 1997, Metzger and Herrel, 2005)** and dinosaurs **(Barrett, 2014)**. With the absence of hydrodynamic constraints, the oreinirostral skulls of Mesozoic terrestrial crocodyliformes are expected to be more efficient at resisting feeding-induced loads than their extant and extinct semi-aquatic platyrostral counterparts.

While the function of platyrostral and oreinirostral skulls has been compared in theoretical skull shapes **(Rayfield and Milner, 2008)** and between crocodilians and odontocetes **(McCurry et al., 2017a)**, for example, a comparative comparison between extant platyrostral forms and extinct oreinirostral taxa is lacking. Here we address this by comparing the ability of oreinirostral and platyrostral skulls of living and extinct crocodyliform taxa to resist feeding induced loads. We hypothesise that the absence of hydrodynamic constraint confers greater stress resistance to the oreinirostral terrestrial taxa. We test this by evaluating the comparative biomechanical performance of the skulls of the Mesozoic terrestrial and extant semi-aquatic taxa. We predict that if oreinirostral morphologies were better suited for resisting feeding induced loads they will display lower magnitudes of feeding induced stress when normalised for size. We also predict that oreinirostral taxa will exhibit increased muscular efficiency, allowing them to produce higher bite forces relative to muscle input.

## Institutional abbreviations

MPMA, Museu de Paleontologia de Monte Alto, São Paulo, Brazil; OUVC, Ohio University Vertebrate Collections, Ohio, U.S.A; OUNHM, Oxford University Natural History Museum, Oxford, U.K; NHMUK, Natural History Museum, London, U.K.

## Materials and Methods

### Specimens

Three extinct crocodyliform and three extant crocodilian taxa, covering a range of skull shapes were chosen for a comparative approach (Figure 1). Three extinct terrestrial crocodyliformes from the Cretaceous Adamantina Formation of the Bauru Basin in Brazil were analysed: *Baurusuchus salgadoensis* (MPMA-62-0001/02), a baurusuchid; *Montealtosuchus arrudacamposi* (MPMA-16-0007/04), a peirosaurid, and *Caipirasuchus paulistanus* (MPMA-67-0001/00), a sphagesaurid. All three specimens possess oreinirostral skulls, with theropod-like lateral compression in their snouts **(Carvalho et al., 2005, 2007, Iori and Carvalho, 2011)**. Extant crocodilians chosen were the highly platyrostral American alligator, *Alligator mississippiensis* (OUVC 9761); the Nile crocodile, *Crocodylus niloticus* (OUNHM 13306); and the brevirostrine Cuvier’s dwarf caiman, *Paleosuchus palpebrosus* (OUNHM 1451). Specimens were scanned using medical and industrial scanners at various locations, with voxel sizes varying between 0.63 mm x 0.63 mm x 0.63 mm (low resolution) for *Montealtosuchus* and 0.099 mm x 0.099 mm x 0.099 mm (high resolution) for *Paleosuchus* (Table S1). This variation in scan resolution is expected have very little influence on the results for such comparative studies where the specimens are many orders of magnitude greater in size than the scan resolution **(McCurry et al., 2015)**.

**Figure 1.**
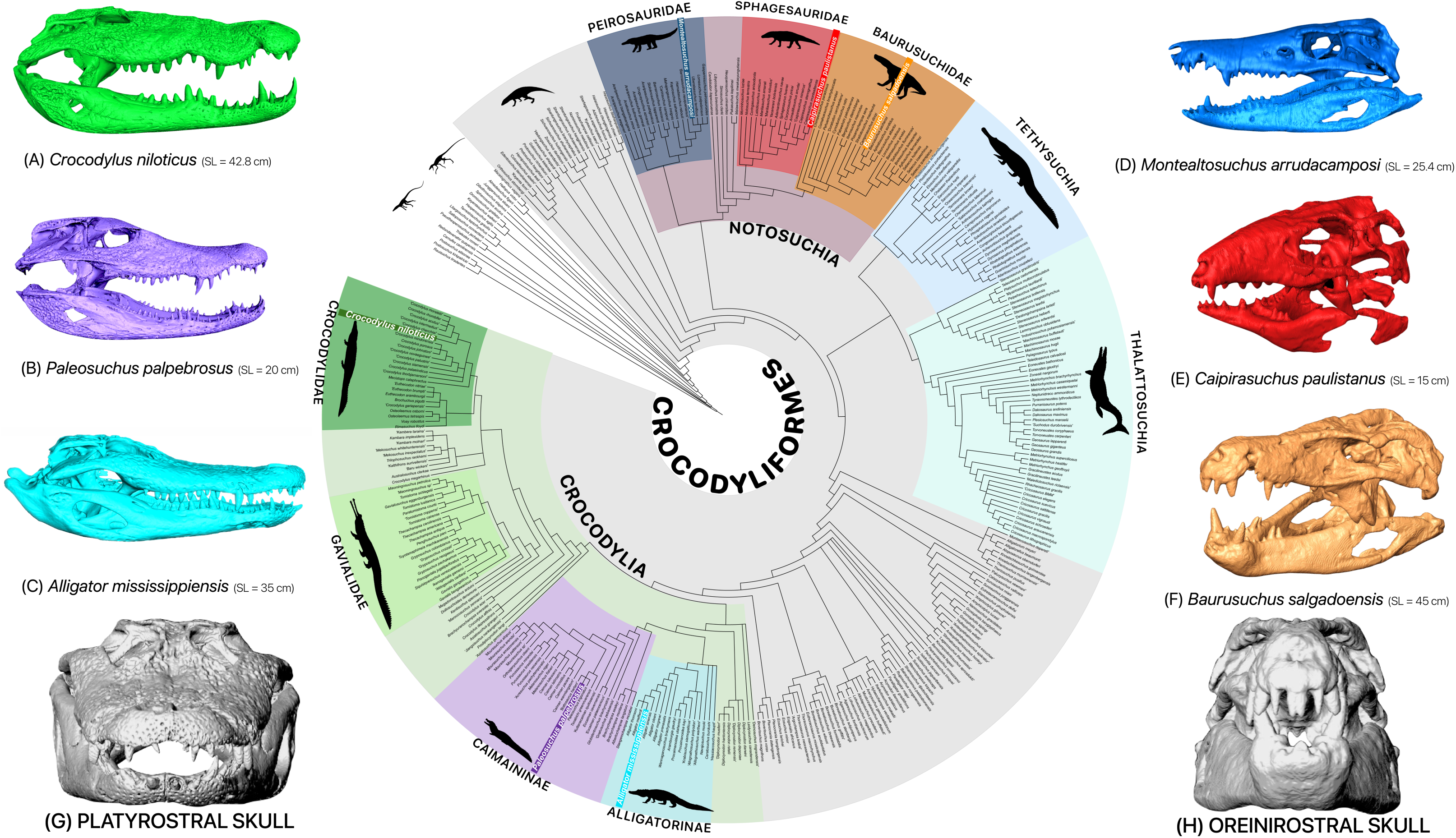
Digitally segmented models of taxa used in this study, along with their phylogenetic positions within Crocodylomorpha. Extant semi-aquatic platyrostral crocodilians: (A) *Crocodylus niloticus* (OUNHM 13306), (B) *Paleosuchus palpebrosus* (OUNHM 1451), and (C) *Alligator mississippiensis* (OUVC 9761). Extinct terrestrial oreinirostral notosuchians: (D) *Montealtosuchus arrudacamposi* (MPMA-16-0007/04), (E) C*aipirasuchus paulistanus* (MPMA-67-0001/00), and (F) *Baurusuchus salgadoensis* (MPMA-62-0001/02). (G, H) Anterior view of platyrostral morphology in *Alligator mississippiensis* and oreinirostral morphology in *Baurusuchus salgadoensis*. SL denotes skull length. Tree data from Stubbs et al. 2021. Silhouettes from PhyloPic (http://phylopic.org; see Table S4 for attributions).

### Model Creation

The computed tomography datasets were visualised and digitally segmented using Avizo 9.4.0 (Thermofisher Scientific) to generate three-dimensional surface models required for finite element modeling. The datasets were segmented into cranium and mandible using Avizo’s segmentation editor (Figure 1A-F). Due to limitations in scan resolution and preservation quality, enamel, dentine and tooth root could not be reliably isolated from bone in the fossil scans. Therefore, to keep model anatomy comparable, all teeth were given properties of bone across all taxa. Herbst et al. (2021) demonstrated that simplifying dental materials does not impact stress distribution comparisons among distantly related taxa, it can alter bite force estimates depending on the specific material properties assigned. Treating teeth as bone may lead to under- or overestimation of bite forces in FE models, depending on the enamel thickness and general tooth structure **(Herbst et al., 2021)**. Given both the lower resolution of fossil scans and the variation in enamel structure between taxa and tooth types, our adopted uniform approach likely yields conservative estimates for bite force ranges. Although sutures can impact the mechanical performance of a skull by decreasing the strain in the surrounding bones **(Jaslow and Biewener, 1995, Rayfield, 2005, Reed et al., 2011)**, they were also not modeled here as this information was not preserved in the fossil taxa. During segmentation, the fossil models were repaired by filling in the cracks and fixing missing parts of bones by use of interpolation **(Lautenschlager, 2016)**. The right side of the cranium of *Caipirasuchus* (MPMA 67-0001/00) was missing the quadrate, parts of the lateral temporal fenestra, and the region enclosing the otic chamber. The model was reconstructed by mirroring and merging the left side across the mid-sagittal plane, using well-known digital restoration techniques **(Lautenschlager, 2016)**. The left side of *Baurusuchus* (MPMA-62-0001/02) is relatively well preserved and complete, while the right side is taphonomically deformed and incomplete **(Carvalho et al., 2005)**. However, as the right side of the cranium was plastically deformed into the left side, it could not be readily restored by mirroring, as an exact mid-sagittal plane could not be established (Figure S1 a-d). The model was imported into the geometric morphometrics (GMM) software LANDMARK **(Wiley et al., 2007)**, and by applying a series of corresponding landmark primitives to either side of the mid-sagittal axis (Supplementary methods), the cranium was retrodeformed (Figure S1 e-h). The resultant model was less plastically distorted, and was used to reflect the components of the left side. The *Montealtosuchus* (MPMA-16-0007/04) skull model was well preserved and did not require any restoration.

### Digital Muscle Reconstructions

In order to estimate forces generated by muscle contraction, jaw adductor muscles were digitally reconstructed in *Baurusuchus,* following previous three-dimensional digital muscle reconstruction methods **(Lautenschlager, 2013, Lautenschlager, 2015)**. Osteological correlates and surface features such as muscle scars and depressions were used to identify sites of muscle origin and insertion in the 3D models. The surfaces of muscle attachment and the dimensions were studied from literature on crocodile jaw musculature **(Iordansky, 1964, 2000, Holliday et al., 2013)** to assist in the reconstructions. The eight muscles modeled were the adductor mandibulae posterior (m.AMP); the adductor mandibulae externus medialis (m.AMEM), superficialis (m.AMES) and profundus (m.AMEP); the pseudotemporalis superficialis (m.PSTs) and profundus (m.PSTp); and the pterygoideus dorsalis (m.PTd) and ventralis (m.PTv) **(Holliday and Witmer, 2007)**. In numerous sauropsids, including extant crocodilians and fossil crocodyliformes, the intramandibularis muscle (m.IM) and the pseudotemporalis superficialis (m.PSTs) muscles pass through the cartilago transiliens, a fibrocartilaginous structure that facilitates muscle attachment, force transmission and redirection **(Goessling et al., 2008, Tsai and Holliday, 2011)**. However, due to the lack of osteological correlates in the fossil specimens, this structure and other tendinous attachments were not modeled separately. The m.PSTs and m.IM were instead reconstructed together as a single muscle complex (m.PSTs), which represents the intramuscular continuum found in crocodilians **(Tsai and Holliday, 2011)**, and does not affect muscle volume estimates. To create the digital muscles, the identified origin and insertion sites were connected by ten simple cylindrical beams per muscle using the labeling and interpolation functions in Avizo. It was ensured that these beams did not traverse through the bone or each other. The muscles were then increased in size uniformly until they were in contact with each other or the bone, thus producing a ‘fleshed’ model. The individual muscle volumes were then calculated using the ‘Material Statistics’ function in Avizo. While previous FE-studies **(Walmsley et al., 2013, McCurry et al., 2017a)** have generally adapted a reptile version of Thomason’s ‘dry skull method’ for mammals **(Thomason, 1991)**, here physiological cross-sectional area (PCSA) was estimated directly by dividing the virtual muscle volumes (*Vm*) by the average fibre length (*Fl*) of the reconstructed muscles. The measure tool in Avizo was used to calculate the mean muscle length using ten estimates between the points of origin and insertion. Most recent studies have assumed muscle fibre length as equal to muscle length (M*l*). Here, fibre lengths were calculated as being a third of the total muscle length to account for the fact that fibres frequently do not stretch along the entire length of the muscle **(Bates and Falkingham, 2018)**. This value, determined using reduced major axis regression by Bates & Falkingham (2018) from published datasets on muscle architecture of 1,100 muscles from extant terrestrial vertebrates, is also consistent with direct measurements from a juvenile *Alligator* where F*l*:M*l* ranges from 0.28–0.9 (mean 0.47) **(Porro et al., 2011)**. The estimated PCSAs were then multiplied with a value for isometric muscle stress (**σ**) of 0.3 N mm^-2^ to obtain the muscle forces **(Thomason, 1991)**.

Sellers et al. **(2022)** noted conservatism among jaw muscle proportions in crocodyliformes. Accordingly, each individually reconstructed muscle in *Baurusuchus* was assumed to occupy the same proportion in other taxa and was modelled on to the corresponding muscle attachment sites (Figure S2). The resultant individual muscle forces were scaled in the other taxa using scaling ratios of the surface area of the models, and the calculated surface area of the muscle attachment sites to fit the available volume of the skull, as measured in Avizo [Equation 1].

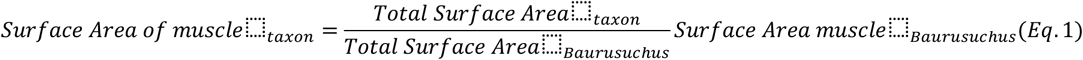

The force to surface area ratio was kept constant across all models in order to facilitate a comparative analysis of rostral morphology **(Dumont et al., 2009)**.

### Finite Element Modelling

Surface models of each cranium were imported from Avizo into HyperMesh v14 (Altair Engineering) for generating 3D meshes. Meshes were downsampled using the ‘Shrink wrap’ function in Hypermesh to reduce complexity and comprised between 725,000 and 15,000,000 tetrahedral elements (Table S2). In order to test for variations in the results caused by the number of surface elements, mesh convergence tests were performed in all models by varying element sizes between 0.5 mm to 4 mm **(McCurry et al., 2015)**. An element size of 0.9 mm was chosen for all models, as mesh convergence was achieved between 0.7 mm and 1 mm.

Bone was treated as elastic and isotropic. Material properties measured *in vivo* from *Alligator* mandibular cortical bone (Young’s modulus (E) = 15 GPa and Poisson’s ratio (***v***) = 0.29) **(Zapata et al., 2010, Porro et al., 2011)** were applied to the model. The FE models were restrained in all six degrees of freedom at the quadrate condyle using thirty-five nodes on the right side, and in the anteroposterior and dorsoventral axes on the left side. This simulates reaction between the skull and mandible, and prevents overconstraint of the models by allowing the skull to flex mediolaterally at the jaw joints (Figure S3). Two sets of functionally comparable tooth positions were loaded, rather than anatomically comparable positions. These were termed ‘caniniform’ and ‘molariform’ **(Erickson et al., 2012)**, and were constrained in the dorsoventral direction using ten nodes per side. The calculated muscle forces were then applied to the models as equally distributed loads at points of muscle origination and insertion, following the direction of insertion, with fifty to one hundred nodes per muscle. To analyse feeding behavior in the crocodyliformes, we tested bilateral biting at the (i) caniniform and (ii) molariform teeth, (iii) bilateral pull-back action at the caniniform teeth, and (iv) unilateral biting at the caniniform teeth. The first three simulations represent the predicted feeding behavior in the extinct forms **(Busbey, 1995)**, while unilateral caniniform biting represents rolling behavior, a commonly used form of prey capture in the extant forms. For the pull-back action, models were constrained on the right quadrate condyle in the dorsoventral and mediolateral axes, and on the left in the dorsoventral axis. An additional constraint was applied at the occipital condyle in the anteroposterior axis using ten nodes to simulate reaction forces at the neck. Loads were applied to one node on either side, on the distal side of the caniniform teeth, based on the bite forces generated during bilateral caniniform biting in the vertical direction (Table S3).

Models were imported into and solved using Abaqus FEA 6.14 (Dassault Systèmes Simulia; https://www.3ds.com). Biomechanical performance was primarily evaluated by comparing the contour plots of the von Mises stresses, which represents tensile and compressive stresses and determines where a material is likely to yield. Also measured were metrics for maximum and minimum principal stresses and strains, which represent tensile and compressive loads respectively. Median von Mises stress values were computed using code in R v3.4.2 **(Team, 2013)**. The top 5% values were excluded to remove the influence of high stress artefacts occurring at the condyle and bite point constraint locations **(Walmsley et al., 2013)**. Bite forces for each taxon were estimated by measuring the reaction forces generated by the FE models. Subsequently, bite forces were divided by the input muscle forces to calculate muscle efficiency.

## Results

### Adductor Musculature and Muscle Forces

The location and spatial arrangement of adductor muscle anatomy in *Baurusuchus* shares similarities with those of extant crocodilians, but the muscles are elongated and less medially aligned (Figure 2 a-c; S4-S5). A key difference in the musculature is found with the pterygoideus group (Figure S5 c-d). The pterygoid flanges in *Baurusuchus* are concave and vertically oriented **(Carvalho et al., 2005)** (Figure S6),allowing for an expansive area of m.PT attachment, and contributing to larger pterygoideus muscle volumes (Figure 2d). In *Baurusuchus*, osteological evidence suggests that the lateral surface of the angular serves as an insertion site for the m.PTv. This trait is synapomorphic in all extant crocodilians, but is absent in numerous crocodyliformes, including several sebecosuchians, peirosaurids such as *Montealtosuchus*, and other derived notosuchians **(Sellers et al., 2022)**. PCSA estimates show that the m.PT, m.AMP and m.PSTs occupy the largest volumes and account for the majority of muscle force (Figure 2e).

**Figure 2.**
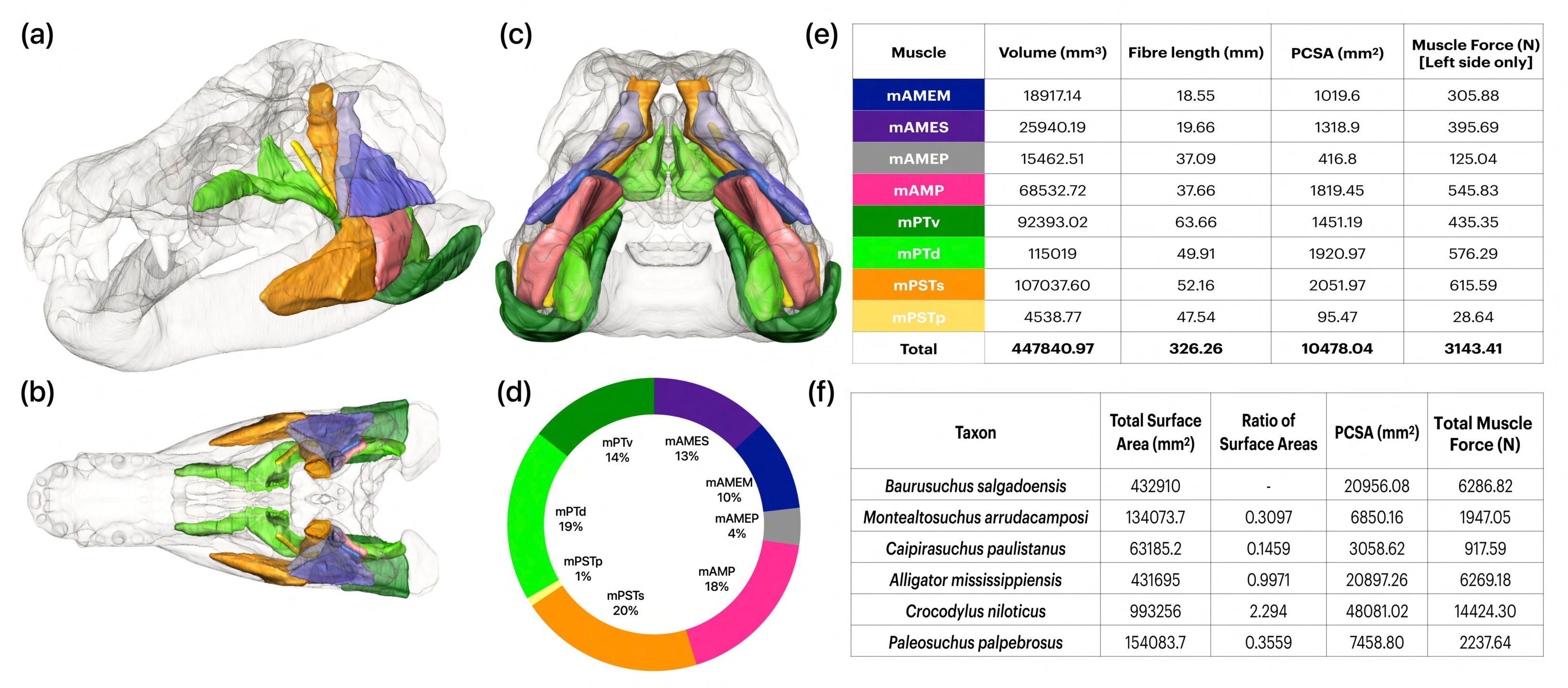
Digitally reconstructed adductor muscles in *Baurusuchus salgadoensis* in (a) dorsal, (b) lateral and (c) ventral views; (d) Chart depicting the proportions of input muscle forces; (e) Table of volumes, fibre lengths, physiological cross-sectional areas and muscle forces for one side of the skull in *B.salgadoensis*; (f) Table of total scaled muscle forces for the entire skull for all other taxa, assumed to occupy the same proportions as in (d).

### Finite Element Analysis (FEA)

Our simulations show that platyrostral taxa experience higher magnitudes of stress than the oreinirostral taxa, in all feeding scenarios. During caniniform bilateral biting, high levels of stress were localised at the pterygoid flanges, skull roof, palate, and the biting teeth (Figure 3). The ventral surfaces of the pterygoid flange and ectopterygoid show relatively higher magnitudes of maximum principal stresses (Figure S7). In contrast, the quadrates, margins of the orbit and maxilla, and the dorsal portion of the pterygoid and epipterygoid experience minimum principal stresses (Figure S8). As with the von Mises stresses, the magnitudes of these metrics are markedly higher in the extant forms than the extinct forms. During the pull-back loading, higher concentrations of stress were observed in all taxa in the quadrate, squamosal, quadratojugal, and jugal. In the platyrostral taxa, this also extends to the maxilla (Figure 3 d-f). During bilateral biting at the molariform tooth, the crania were under lower stress than during caniniform biting and an increase in the total stress can be observed towards the posterior region where the muscle forces and boundary conditions are modelled. During unilateral biting at the caniniform tooth, stress patterns were much higher in magnitude on the working side and relatively low on the balancing side. Areas experiencing comparatively high stress levels include the premaxilla, the maxilla, and the jugal in the extant forms. Patterns of stress and strain distributions (Figure S9) remain broadly similar across the different taxa, irrespective of the loading scenario. Median stress values follow the patterns established by the contour plots of the FE models, with *Caipirasuchus* experiencing the lowest median stress, followed by *Baurusuchus*, whereas *Paleosuchus* exhibits the highest stress (Figure 4a).

**Figure 3.**
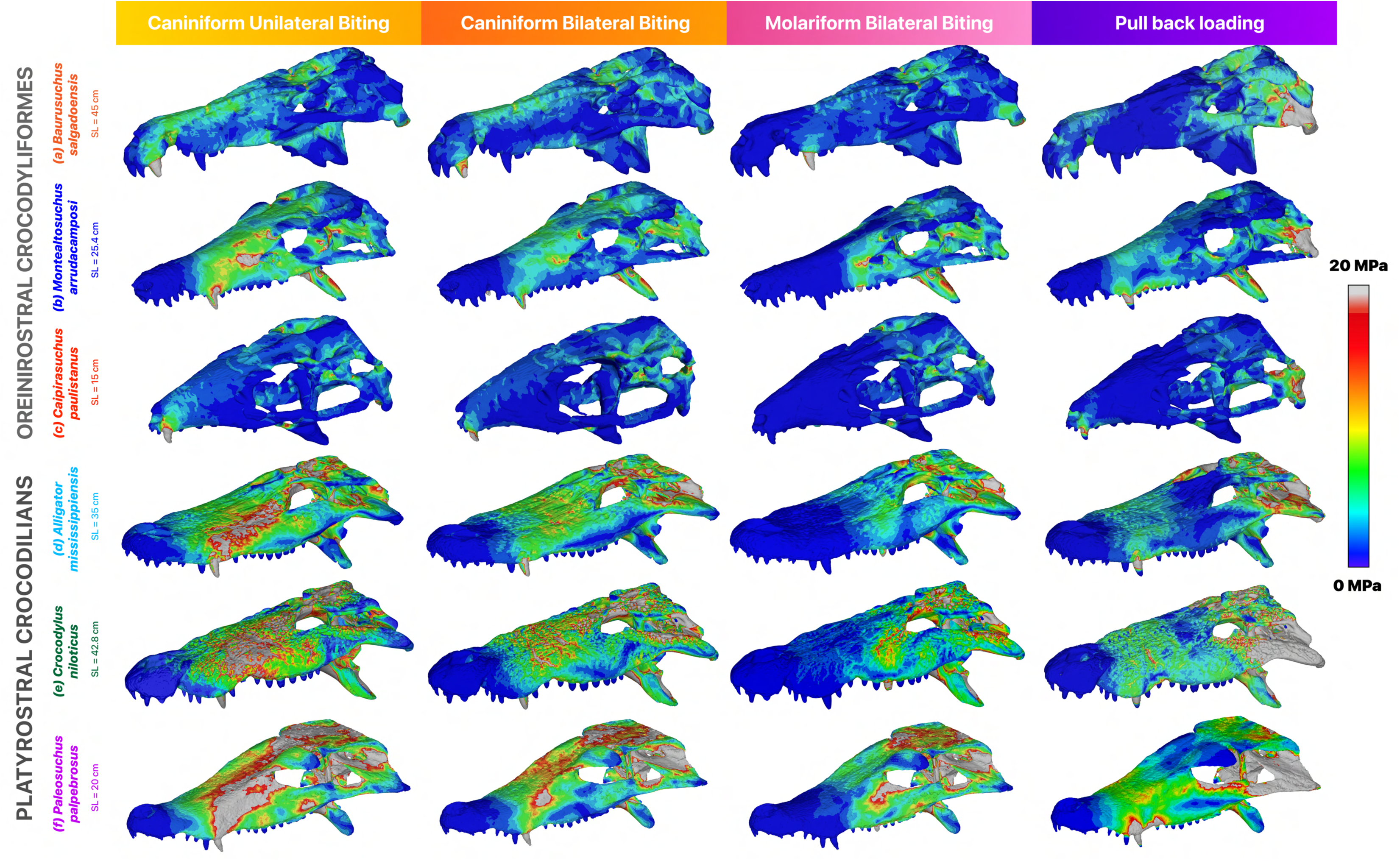
From top to bottom: von Mises stress distribution in the cranium during various feeding scenarios for (a) *Baurusuchus salgadoensis,* (b) *Montealtosuchus arrudacamposi,* (c) C*aipirasuchus paulistanus,* (d) *Alligator mississippiensis,* (e) *Crocodylus niloticus* and (f) *Paleosuchus palpebrosus.* SL denotes skull length. Areas in grey depict stresses greater than 20 MPa.

**Figure 4.**
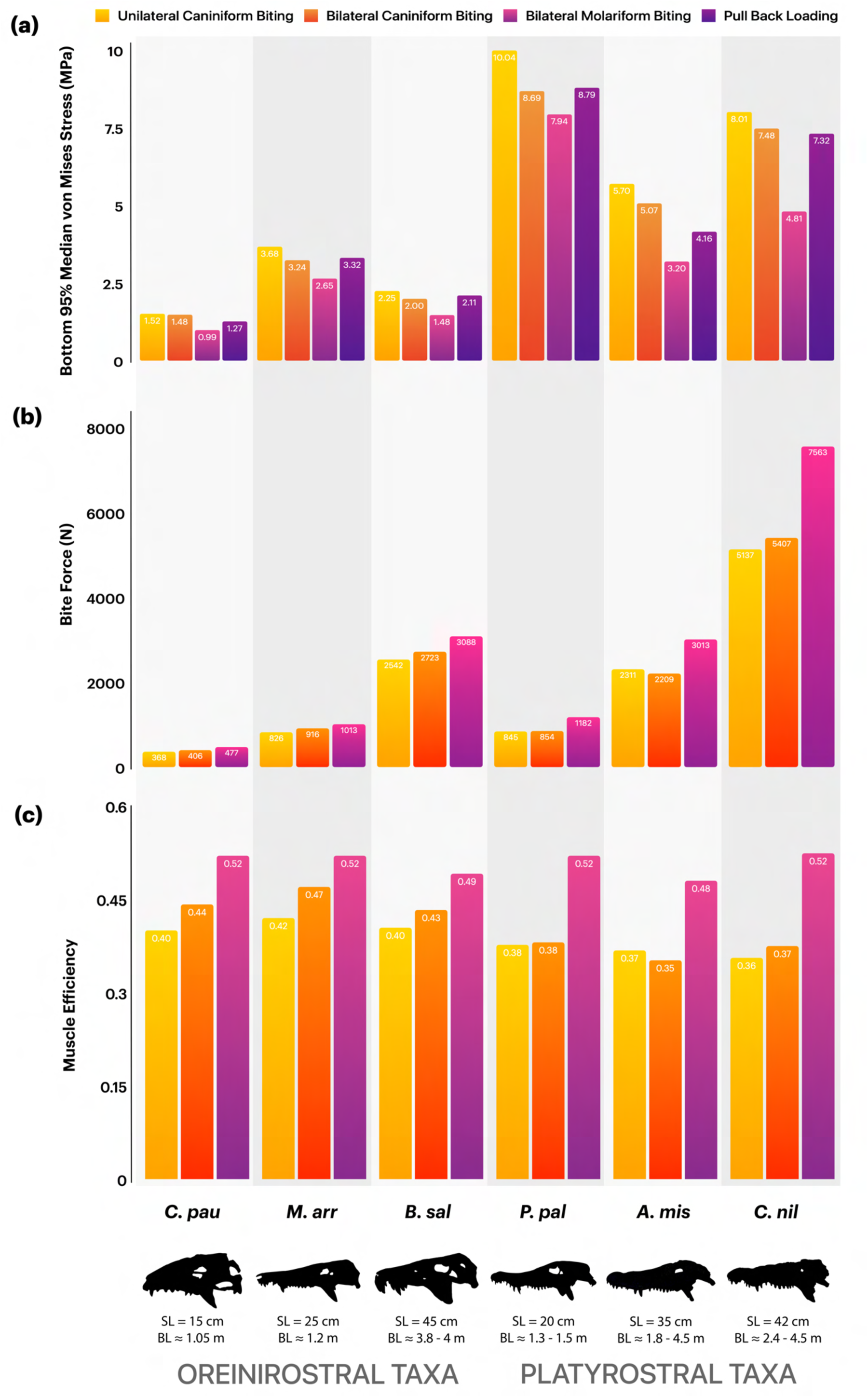
From top to bottom: (a) 95% median von Mises stress values, (b) bite forces, and (c) muscle efficiency estimates for (left to right) *Caipirasuchus paulistanus, Montealtosuchus arrudacamposi, Baurusuchus salgadoensis, Alligator mississippiensis, Crocodylus niloticus* and *Paleosuchus palpebrosus* under different loading regimes. SL and BL denote measured skull lengths and estimated body lengths respectively.

### Bite Forces and Muscular Efficiency

Bite force measures increased from the anterior to the posterior region, due to the reduction of the distance between the bite point and jaw joint (Figure 4b). *Caipirasuchus* was found to have the lowest bite forces, ranging from 368 to 477 N, while *Baurusuchus* and the *Alligator* show relatively similar bite forces from 2,542 to 3,088 N, and 2,311 to 3,013 N, respectively. *Crocodylus* has the largest bite forces from 5,137 N to 7,563 N. Bite force values for the extant taxa are consistent with those that were reported from *in vivo* tests in living crocodilians of similar sizes **(Erickson et al., 2012, Sellers et al., 2017)**. Oreinirostral taxa transfer muscle forces into bite forces with increased efficiency compared to platyrostral taxa during unilateral and bilateral caniniform biting (Figure 4c). Platyrostral taxa show similar muscular efficiency between unilateral and bilateral caniniform biting. During bilateral molariform biting, all taxa possess similar muscle efficiencies, due to shortened lever arms.

## Discussion

### Biomechanical and functional adaptations in oreinirostral crocodyliformes

Our primary aim was to compare the ability of oreinirostral and platyrostral skulls of extant and extinct crocodyliformes to resist feeding induced loads, hypothesising that the absence of hydrodynamic constraints in terrestrial oreinirostral taxa confers greater stress resistance. The results from the finite element analysis confirm our initial predictions. When muscle loads are scaled to skull surface area (normalising for size), extinct terrestrial crocodyliformes with oreinirostral morphologies consistently produce lower stress values during feeding than extant platyrostral crocodilians. The following inferences assume that lower stress and strain magnitudes and greater dispersion of stresses for models loaded under similar constraints, implies increased structural strength before failure **(Rayfield et al., 2001, Rayfield, 2005, Fastnacht et al., 2002, Preuschoft and Witzel, 2002)**.

Busbey (1995) hypothesised that platyrostral morphologies were sub-optimal compared to oreinirostral ones, and that the evolution of platyrostry was linked to better resisting torsional loads, developed as a functional trade-off between hydrodynamic constraints and feeding-induced loads. The results of this study find partial support for Busbey’s hypothesis, in that platyrostral morphologies are biomechanically sub-optimal at resisting feeding-induced loads, compared to oreinirostral morphologies. However, we find no evidence that the evolution of platyrostry was linked to better resisting torsional loads, consistent with previous studies **(Metzger et al., 2005, McHenry et al., 2006, Nieto et al., 2022)**. Our study also finds support in Busbey’s prediction that sebecosuchians and other larger terrestrial oreinirostral crocodyliformes likely fed in a manner similar to the Komodo dragon (*Varanus komodoensis)*, employing a “slice-and-pull-back" action **(Busbey, 1995)**. This type of feeding mechanism generates dorsoventral compressive loads, while minimising torque, influencing both feeding mechanics and overall cranial function **(Carvalho et al., 2005)**. Previous studies of *Varanus* ecology and skull mechanics demonstrate that this feeding strategy distributes stress efficiently throughout the skull, minimising localised strain **(Auffenberg, 1981, Moreno et al., 2008)**. These minimisations are also replicated by the extinct taxa in this study under extrinsic pull-back loading conditions (Figures 3), with oreinirostral morphologies exhibiting lower stress values and greater structural efficiency in handling such movements, compared to platyrostral forms. Our pull-back results are also consistent with those of FEA studies in other notosuchians such as *Baurusuchus pachecoi* **(Montefeltro et al., 2019)**, and *Araripesuchus gomesii* **(Nieto et al., 2022)**, reinforcing the prevalence of this type of feeding behavior in oreinirostral terrestrial crocodyliformes. The presence of antorbital fenestrae in *Caipirasuchus* and *Montealtosuchus* does not present as a region of weakness in the skull, as previously assumed **(Rayfield and Milner, 2008)**, due to the presence of an ossified secondary palate: a feature that is thought to have evolved repeatedly in crocodylomorphs as a response to different evolutionary pressures, such as diet (herbivory, hypercarnivory) or morphology (platyrostry, longirostry) **(Dollman and Choiniere, 2022)**.

We also hypothesised that oreinirostral forms would exhibit increased muscular efficiency. Consistent with previous studies on both crocodilians and other lepidosaurs, our bite force results scale isometrically with both body length and head size (Figure 4b) **(Erickson et al., 2012, Isip et al., 2022)**. Deeming **(2022)** highlights that increases in jaw muscle mass may drive the evolution of larger heads (or conversely, that larger heads may accommodate more jaw musculature), and thus producing higher bite forces, while also emphasising the importance of mechanical advantage (or muscle efficiency). Mechanical advantage seemingly remains constant across reptiles of different sizes **(Deeming, 2022)**, suggesting that variations in bite force are primarily driven by muscle mass and head dimensions rather than changes in moment arm ratios. Hence, the relationship between skull shape, skull size, jaw musculature, and bite force may be central to understanding the cranial evolution of crocodyliformes. In *Baurusuchus salgadoensis*, cranial modifications, such as an elevated snout, elongated quadrate, and ventrally deflected pterygoid flanges create larger muscle attachment areas and optimise muscle pull direction, strengthening the skull (Figure 3a). This also allows for longer muscle fibers (Figure S4-S5), which directly influences muscle forces. Several muscle attachment sites, including the supratemporal fossa and the mandibular fossa, where the m.AMEP and the m.PSTs attach, respectively, are significantly larger in the extinct crocodyliformes in this study compared to extant crocodilians. Many studies have attributed the smooth surface of the supratemporal fossa in archosaurs as a muscle attachment site. In our reconstructions for *Baurusuchus* we did not attach the aforementioned muscles to this region, which serves as an attachment for other vascular tissues. Instead, the m.AMEP and m.PSTs attach to the dorsotemporal fossa **(Holliday et al., 2020)**. Additionally, the anterior surface of the articular, which serves as the insertion site for the m.AMP, m.PTd, and m.PTv, is concave in the oreinirostral taxa and flatter in extant taxa, and also provides an expanded area for muscle attachment, (a feature also observed in Sebecus **(Molnar, 2012)**). Adductor muscle orientation in oreinirostral skulls increased mechanical efficiency for terrestrial predation **(Busbey, 1989, Stubbs et al., 2013)**, allowing terrestrial crocodyliformes to be biomechanically optimised for bite force efficiency rather than absolute force magnitude.

### Functional adaptations in the skull for feeding ecology and diversity in notosuchians

Oreinirostral crocodyliformes evolved robust cranial geometries, enabling efficient force distribution adapted for different modes of feeding and diets. Although high rates of morphological evolution have been observed within both Notosuchia and Crocodilia, there is high variability among notosuchians in regions such as the pterygoid, ectopterygoid, jugal, quadratojugal, and squamosal **(Felice et al., 2021)**, which are biomechanically relevant and show greater resistances to feeding induced loads in the oreinirostral taxa in our study. This variability probably allowed notosuchians to explore various ecological niches **(Ősi, 2014, Melstrom and Irmis, 2019)**. This contrasts with the relatively conserved cranial architecture of semi-aquatic crocodilians, where stabilising selection maintained platyrostry despite developmental shifts in snout elongation. Most cranial elements remain unchanged in extant species, except for the quadrate and pterygoid flanges, which aid in producing large bite forces **(Felice et al., 2021)**.

Among sphagesaurids, *Caipirasuchus paulistanus*possessed the tallest skull and a proportionately large supratemporal fossa, which may have enhanced its ability to resist feeding induced loads **(Iori and Carvalho, 2011)**. Despite its small size (∼1.05 m) **(Iori et al., 2016)**, *Caipirasuchus* exhibits cranial morphology well optimised for withstanding feeding induced loads, and consistently shows the lowest stress values among all taxa analysed in this study (Figure 3c). The unique bulbous dentition observed in *Caipirasuchus* and other sphagesaurids such as *Morrinhosuchus,* have led to possible interpretations of herbivory or omnivory in certain notosuchians **(Nobre et al., 2008, Ősi, 2014, Iori and Carvalho, 2018)**. In addition, *Caipirasuchus* exhibits complex tooth replacement patterns **(Borsoni and Carvalho, 2024)**, and sphagesaurids are noted to have increased enamel thickness over baurusuchids **(Ricart et al., 2021)**, indicating a herbivorous diet. Thicker enamel in herbivorous and omnivorous taxa likely enhances wear resistance from extensive food processing, while carnivorous taxa exhibit thinner enamel, as seen in other crocodilians and lepidosaurs **(Kay, 2017, Sellers et al., 2019)**. These adaptations may have aided *Caipirasuchus* during propalinal jaw movements, which are anterior-posterior sliding motions facilitated by specialised jaw joint morphology; as well as other complex mastication mechanisms **(Marinho and Carvalho, 2007, Iori and Carvalho, 2011, Iori and Carvalho, 2018)**. Such mechanisms were once thought to be rare among crocodyliformes, but Melstrom and Irmis **(2019)** report multiple independent origins of herbivorous adaptations within this group, often exhibiting convergent traits such as complex dentition and robust skull morphologies. Though they did not sample *Caipirasuchus*, these features are evident in this species, suggesting functional adaptations for herbivory or omnivory. Its size and feeding strategies may have enabled it to exploit new resources, highlighting ecological partitioning within notosuchians.

The cranial design of *Montealtosuchus* allows for efficient dissipation of stress across the skull, with increased muscular efficiency during both unilateral and bilateral biting. Patterns of stress distribution in this taxon are similar to those observed in *Araripesuchus gomesii* **(Nieto et al., 2022)**, a smaller uruguaysuchid interpreted as an omnivore feeding on insects, small vertebrates, and soft plant material **(Sereno and Larsson, 2009, Ősi, 2014)**. However, our bite force calculations (828–1,013 N), and increased muscle efficiency, suggest *Montealtosuchus* could generate strong, controlled bites for crushing and subduing prey rather than for quick snapping. This, combined with its larger size **(∼ 1.2 m, Tavares et al., 2017)**, robust cranial and postcranial anatomy, laterally positioned orbits, semi-upright gait, and light dermal armor **(Carvalho, 2007, Tavares et al., 2015)**, suggests a cursorial predatory lifestyle, ambushing both medium and small-sized prey in the semi-arid floodplain environments of the Adamantina Formation. Cranial morphological similarities between *Montealtosuchus* and early-diverging crocodyliformes like *Caririsuchus camposi* (lower Albian, Brazil) and *Hamadasuchus rebouli* (Albian–Cenomanian, Morocco) suggest that peirosaurids retained ancestral traits adapted for a generalist diet. This conservatism may have allowed *Montealtosuchus* to occupy a mesopredator niche, alongside late-diverging baurusuchids and sebecids that evolved hypercarnivorous adaptations to fill apex predator roles.

Among notosuchians, baurusuchids are a highly derived group, exhibiting several traits consistent with hypercarnivory, including anterior positioning of the external nares, lateral compression of the snout, reduction in the number of teeth, ziphodont dentition, and enlarged caniniform teeth **(Carvalho et al., 2005, Pinheiro et al., 2008)**. *Baurusuchus salgadoensis* evolved highly specialised cranial adaptations that enabled it to thrive as an apex predator in the Late Cretaceous floodplain ecosystems, alongside other medium-sized theropods **(Gasparini et al., 1993, Candeiro et al., 2024)**. Our functional analysis reveals exceptional resistance to stress during feeding induced loads, particularly in pull-back scenarios where stress is localised to the pterygoid flanges, quadrates, and fused nasals, unlike other taxa in this study where stress is more broadly distributed. Additional stress redistribution is observed in the anterior portions of the skull due to the presence of a large notch along which sutures separate the premaxilla and maxilla, also accommodating the enlarged dentary caniniform teeth of the mandible **(Montefeltro et al., 2011)**. This is a unique adaptation among baurusuchids, which also aids the functional performance of *Baurusuchus pacchecoi* **(Montefeltro et al., 2019)**. These adaptations may support Busbey’s hypothesised “slice-and-pull” feeding strategy. While Montefeltro et al. **(2019)** report a weak bite force for the medium-sized *Baurusuchus pachecoi* (∼ 1.7 m), our results indicate that *Baurusuchus salgadoensis* exhibited a relatively powerful bite (2,542 to 3,088 N) for its body size (∼ 3.8 to 4 m; **[84]**). Apart from size differences between the two species, this discrepancy may be explained by the differences in methodological approaches concerning muscle reconstruction and muscle force calculations. We used digital muscle reconstructions for our model, and used the reconstructed volumes to calculate the PCSA, instead of using the surface area of attachment sites to predict muscle force, with muscle length being a third of the fiber length. Bates & Falkingham **(2018)** demonstrate that longer fibre lengths result in smaller PCSAs, resulting in lower muscle forces, subsequently lower bite forces, and vice versa. Our study does otherwise find consensus with Montefeltro et al. **(2019)**, showing clear differences between oreinirostral and platyrostral skulls, and inferences about cranial adaptations in *Baurusuchus* including hyperossification of the secondary palate, and the absence of antorbital fenestrae. Though *Crocodylus niloticus* generates substantially higher bite forces (5,137–7,563 N; Figure 4b) than *Baurusuchus*, it experiences nearly four times higher stress values (Figure 4a), emphasising a trade-off between raw power and structural efficiency. Similarly, although comparable in size to larger alligators, *Baurusuchus* exhibits much lower stresses and an increased muscle efficiency, emphasising the superior performance of oreinirostral skulls for resisting compressive forces. These biomechanical adaptations suggest that *Baurusuchus* probably relied on precision and efficiency to subdue prey, distinguishing it from extant platyrostral crocodilians that depend on brute force to crush and dismember prey. Juvenile baurusuchids, eg. *Pissarrachampsa sera* LPRP/USP 0049 **(Godoy et al., 2018)** exhibit cranial morphologies resembling other adult notosuchians like *Araripesuchus, Mariliasuchus,* and *Montealtosuchus*, suggesting that baurusuchids retained ancestral morphologies early in development before diverging into specialised forms through peramorphic heterochrony **(Godoy et al., 2018)**. Extended growth trajectories may have allowed *Baurusuchus salgadoensis* and *Stratiotosuchus maxhechti* to achieve larger body sizes, and robust skulls optimised for hypercarnivory, while smaller species like *Baurusuchus pachecoi* likely occupied mesopredator niches within crocodyliform-rich ecosystems.

Notosuchian crocodyliformes exemplify how cranial morphology and feeding mechanics diversified in response to ecological opportunity across Gondwanan ecosystems **(Godoy, 2020)**. Their ability to occupy niches ranging from generalist mesopredation (*Montealtosuchus*) to herbivory (*Caipirasuchus*) and hypercarnivory (*Baurusuchus*) highlights how evolutionary flexibility drove their resilience in dynamic environments.

### The role of hydrodynamic constraints in the evolution of skull form in crocodyliformes

The relatively weaker performance of the platyrostral taxa under all loading conditions suggests that this shape is not the most efficient at resisting forces generated during both simple biting and twist feeding (Figure 3 d-f, 4a). This biomechanical limitation reflects an evolutionary trade-off: dorsoventrally compressed snouts of semi-aquatic and marine crocodyliformes likely evolved to reduce drag during aquatic locomotion, prioritising environmental pressures associated with hydrodynamic efficiency over cranial robusticity and feeding performance **(McHenry et al., 2006)**. As a consequence of skull flattening, platyrostral taxa also have medially aligned adductor musculature **(Sellers et al., 2022)**, requiring greater input forces to produce high bite forces, and exhibiting lower muscle efficiency when compared to oreinirostral crocodyliformes. To mitigate elevated stress concentrations during high-force behaviors like the “death roll,” platyrostral crocodilians developed compensatory adaptations, including closure of the antorbital fenestrae, cranial osteoderms, hypertrophied adductor musculature, and reinforced sutures or scarf joints **(Iordansky, 1973, Busbey, 1995, Preuschoft and Witzel, 2002, Metzger et al., 2005, McHenry et al., 2006, Rayfield and Milner, 2008, Gignac and Erickson, 2016)**. These innovations redistributed mechanical stress across the skull, enabling crocodilians to withstand the demands of ambush predation in aquatic environments despite their flattened cranial geometry. In modern crocodilians, the m.PTd, m.PTv, m.AMEP, m.PSTs, and m.IM are hypertrophied, contributing to greater force production. The ‘zwischensehne’, an internal tendon **(Lakjer, 1926)**, optimises the mechanical advantage of these muscles by dynamically modulating the lever arm length to enhance force transmission **(Molnar, 2012)**.

As crocodyliformes explored new ecologies, skull flattening enhanced hydrodynamic performance, facilitating stealthy locomotion and ambush predation in rivers, lakes, and coastal ecosystems **(Busbey, 1995, McHenry et al., 2006)**. Dorsoventral compression of the skull first appeared in Middle Jurassic neosuchians **(Rayfield and Milner, 2008)**, and can be observed in numerous marine thalattosuchians (eg. *Geosaurus*), tethysuchians (eg. *Sarcosuchus*, *Dryosaurus*), goniopholidids (eg. *Goniopholis*, *Siamosuchus*), paralligatorids (eg. *Paralligator*, *Rugosuchus*) and early eusuchians (eg. *Allodaposuchus, Borealosuchus*). Like extant crocodilians, these groups likely offset weaker skull performance through larger body sizes **(Godoy, 2020)**, or specialised feeding strategies like piscivory, often converging on longirostry (e.g. thoracosaurs, thalattosuchians, and gavialoids) **(Ballell et al., 2019)**. Interestingly, *Dakosaurus maximus* and *Dakosaurus andiniensis* are exceptional among marine thalattosuchians for their oreinirostral skulls and ziphodont dentition, displaying adaptations for hypercarnivory and increased torsional resistance over hydrodynamic efficiency, illustrating how functional demands can decouple cranial shape from ecology, even in fully marine lineages **(Young et al., 2010, 2012)**. In contrast to oreinirostral notosuchians, which were not subject to the same ecological limitations, platyrostral taxa exhibit more conserved skull morphologies due to restricted niche opportunities and environmental conditions **(Stockdale and Benton, 2021, Stubbs et al., 2021)**. Low morphological disparity and repeated evolution of platyrostry provides strong evidence for the influence of ecological and functional constraints on skull shape in crocodilians. Despite these constraints, modern crocodilians thrive as apex predators in semi-aquatic environments, and exhibit some of the highest bite forces **(Erickson et al., 2012)**. Exceptional dietary flexibility, observed in both extinct crocodyliformes and modern crocodilians, has been fundamental to their evolutionary adaptability and longevity, enabling them to persist through dramatic environmental upheavals and mass extinction events **(Melstrom et al., 2025)**. The evolutionary success of platyrostral crocodilians exemplifies how functional trade-offs driven by environmental transitions, can be overcome through integrated mechanical adaptations, suggesting that ecological opportunity, rather than morphological optimization, may be a key driver of innovation.

## Conclusion

This study is the first to present a broad set of comparative data between extinct oreinirostral crocodyliformes and extant platyrostral crocodilians using finite element analysis, to test the role of constraints in shaping form and function. Our results demonstrate that oreinirostral taxa exhibit lower feeding induced stresses and have more efficient skulls, supporting the hypothesis that hydrodynamic constraints influence crocodilian skull evolution. Consistent with previous studies, we find partial support in Busbey’s hypothesis, that platyrostry or skull flattening in crocodilians was evolved as a response to the hydrodynamic constraints acting on the skull, and not to resist torsional loads **(McHenry et al., 2006)**. Crocodilians exemplify how functional trade-offs between feeding efficiency and locomotion drive evolutionary innovation during ecological transitions. Similar patterns are evident in other clades, including archosauriformes such as *Proterochampsa nodosa* **(de Simão-Oliveira et al., 2024)**, and *Riojasuchus tenuisceps* **(Taborda et al., 2023)**, where multifunctional structures like skulls show asymmetric trade-offs, which promotes morphological diversity **(Burress and Muñoz, 2022, Tseng et al., 2023, Sansalone et al., 2024)**. Together, these findings underscore the dynamic interplay of constraints and adaptations in shaping evolutionary innovations across diverse taxa, often favoring ecological opportunity over pure functional optimisation.

