## Supplemental Materials for "Constraints and adaptations in crocodyliform skull evolution"

### SUPPLEMENTARY METHODS

#### Retrodeformation Process

The segmented cranial surface of *Baurusuchus salgadoensis* was imported as an STL into LANDMARK Editor (www.idav.ucdavis.edu/research/EvoMorph). Fourty four landmark primitives (twenty two bilateral pairs), were placed, including single points and curves. Landmarks were placed on both sides of the skull at corresponding locations, including the anterior and posterior orbital margins, supratemporal fossae, narial margins, quadrate condyles, dorsal and ventral margins of the pterygoid flanges, posterior palatine and quadrate margins, and medial margins of the squamosal. These landmarks were chosen based on their anatomical symmetry, relative structural robustness, and reduced likelihood of significant displacement due to taphonomic distortion. Bilateral landmarks were placed on homologous regions to provide geometrically consistent reference points for retrodeformation. Midline and centrally located structures, such as the anterior premaxillary margin and occipital region, were included due to their typically lower susceptibility to lateral shearing. Landmarks were preferentially placed on deeper or buttressed cranial regions, such as the palate and braincase floor, which are less prone to plastic deformation, while structurally delicate or taphonomically variable regions, such as the postorbital bars and lateral jugal surfaces, were avoided. This landmark configuration ensured that retrodeformation was guided by anatomical features most likely to reflect the original morphology of the specimen. Six dimension primitives or pairs of points (5, 7, 8, 11, 14 and 17) were chosen to define and constrain the retrodeformation process. By including more than the minimum four pairs, the plane of symmetry was more tightly constrained. The single-axis method was then applied to perform the retrodeformation.

#### SUPPLEMENTARY FIGURES AND TABLES

**Table S1.** Details of scan locations, resolutions and scanner models for all specimens.

| Taxon | Specimen Number | Scanning Location | Scanner Model | Voxel Size |
|---|---|---|---|---|
| *Alligator mississippiensis* | OUVC 9761 | O'Bleness Memorial Hospital, Ohio | General Electric (GE) LightSpeed Ultra | 0.56 x 0.56 x 0.1 |
| *Baurusuchus salgadoensis* | MPMA 62-0001/02 | Unidade de Radiológica Dr. Fabrício Mallouk, Monte Alto SP, Brazil | Siemens Somatom Spirit | 0.625 x 0.625 x 0.8 |
| *Caipirasuchus paulistanus* | MPMA 67-0001/00 | Hospital das Clínicas da Universidade Federal do Triangulo Mineiro | Siemens Somatom Spirit | 0.429 x 0.429 x 0.3 |
| *Crocodylus niloticus* | OUNHM 13306 | University Of Bristol | Nikon XTH225ST | 0.12 x 0.12 x 0.25 |
| *Montealtosuchus arrudacamposi* | MPMA-16-0 007/04 | Instituto de Radiologia – Faculdade de Medicina de São Paulo, USP, Brasil | Discovery CT750 HD | 0.63 x 0.63 x 0.63 |
| *Paleosuchus palpebrosus* | OUNHM 1451 | University Of Bristol | Nikon XTH225ST | 0.099 x 0.099 x 0.099 |

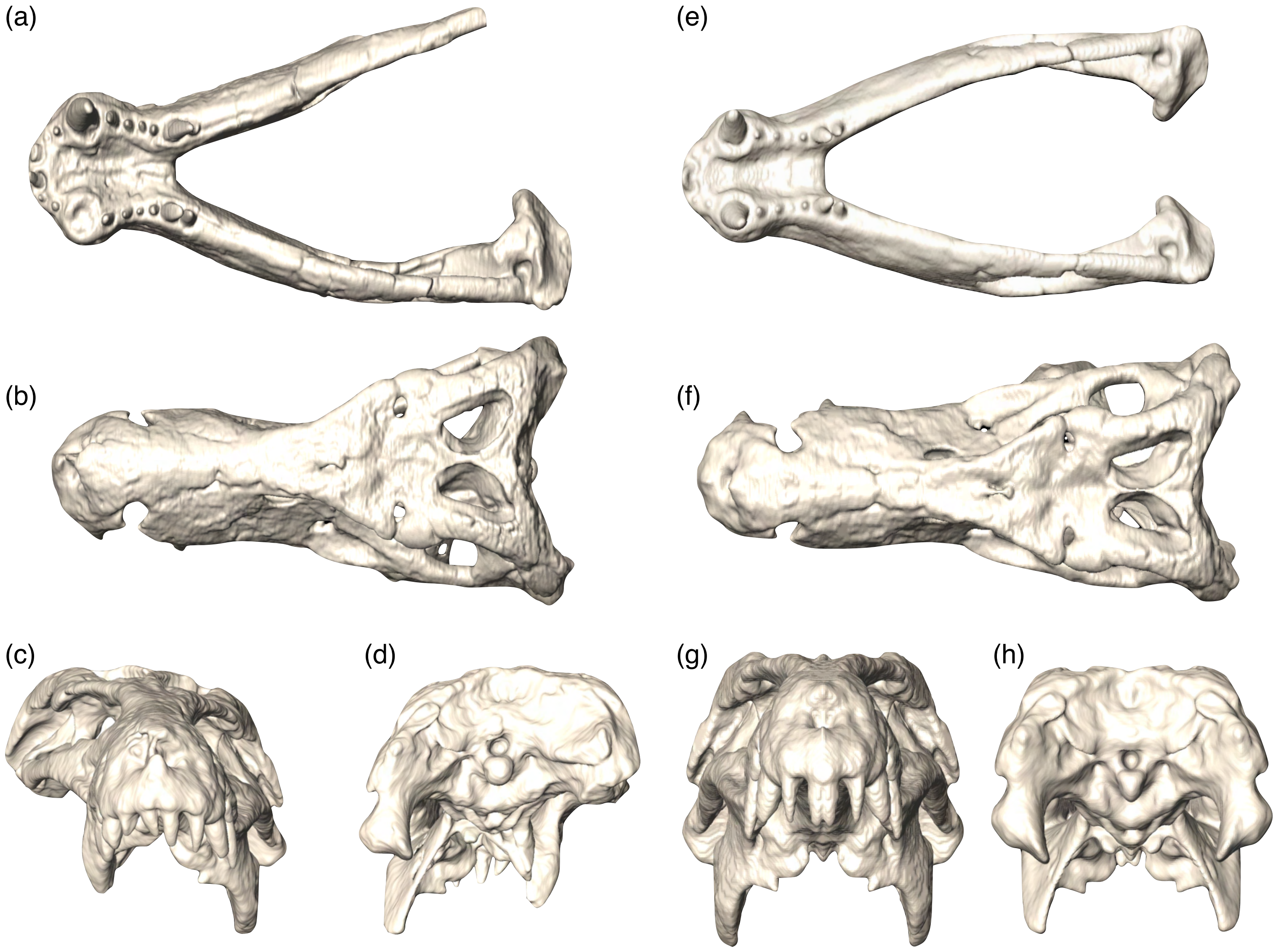


**Figure S1.** Model creation and repairing in *Baurusuchus salgadoensis*. (a-d) Original specimen. (e-h) Restored model. (a) Deformed incomplete mandible. (b-d) Plastically deformed skull in top, front and rear views. (e) Restored mandible using segmentation and mirroring elements of preserved side. (f-h) Restored cranium after retrodeformation and mirroring.

**Table S2.** Details of 3D meshes used for FEA in cranium of all taxa. Note that all FE models are in absolute sizes, and forces were instead scaled to surface area.

| **Taxon** | **Skull Length (in cm)** | **Nodes** | **Elements** | **Element Size (in mm)** |
|---|---|---|---|---|
| *Alligator mississippiensis* | 35 | 1,005,442 | 4,459,374 | 0.9 |
| *Montealtosuchus arrudacamposi* | 25.4 | 448,061 | 2,036,151 | |
| *Baurusuchus salgadoensis* | 45 | 3,096,878 | 14,629,631 | |
| *Caipirasuchus paulistanus* | 15 | 1,490,937 | 6,968,880 | |
| *Crocodylus niloticus* | 42.8 | 2,012,125 | 9,020,310 | |
| *Paleosuchus palpebrosus* | 20 | 176,800 | 725,316 | |

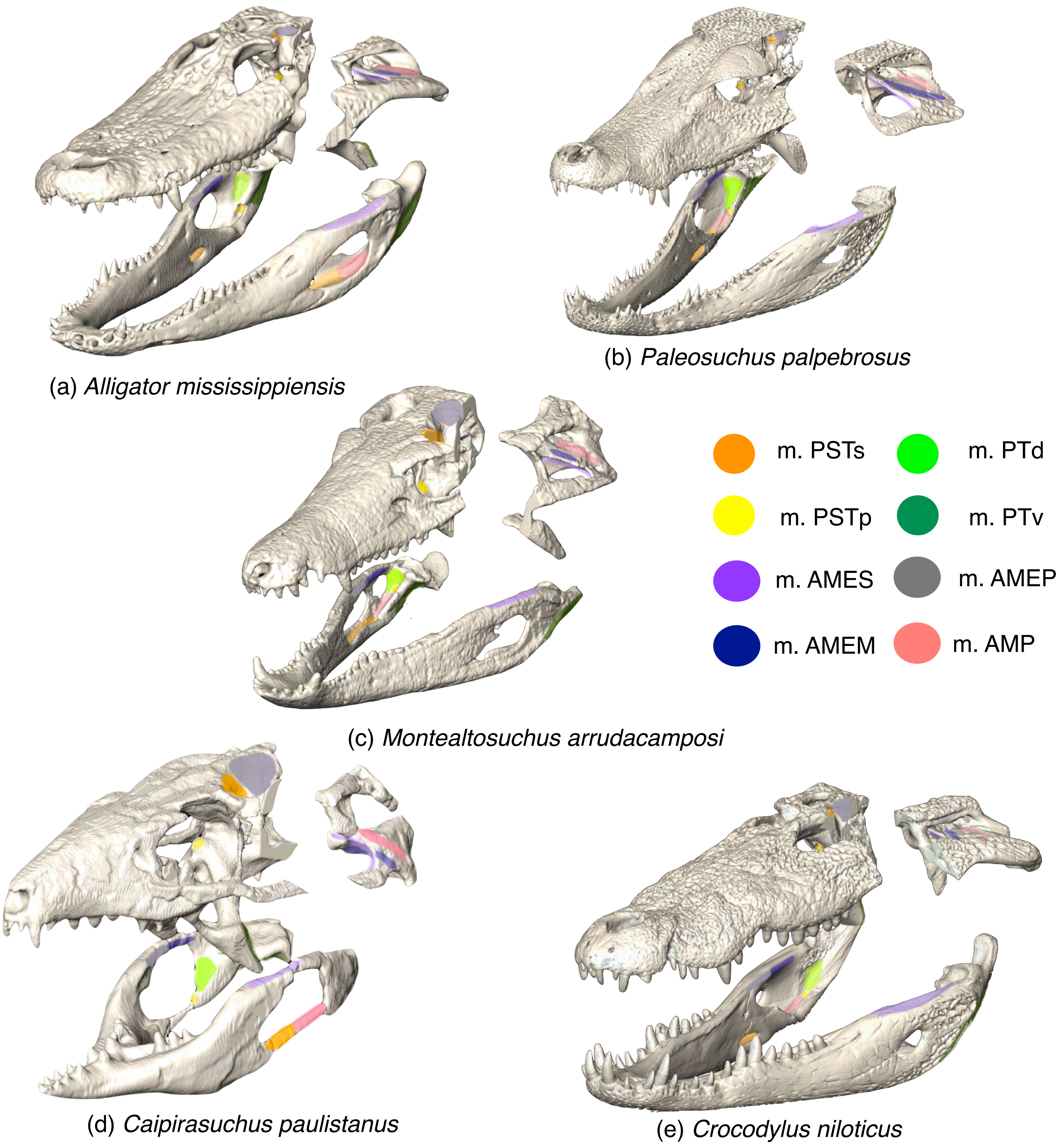


**Figure S2**. Mapping of muscle attachment sites in the taxa (excluding *Baurusuchus*). Note that m.AMES, m.AMP and m.AMEM are found on the inner surfaces of the quadrate and squamosal.

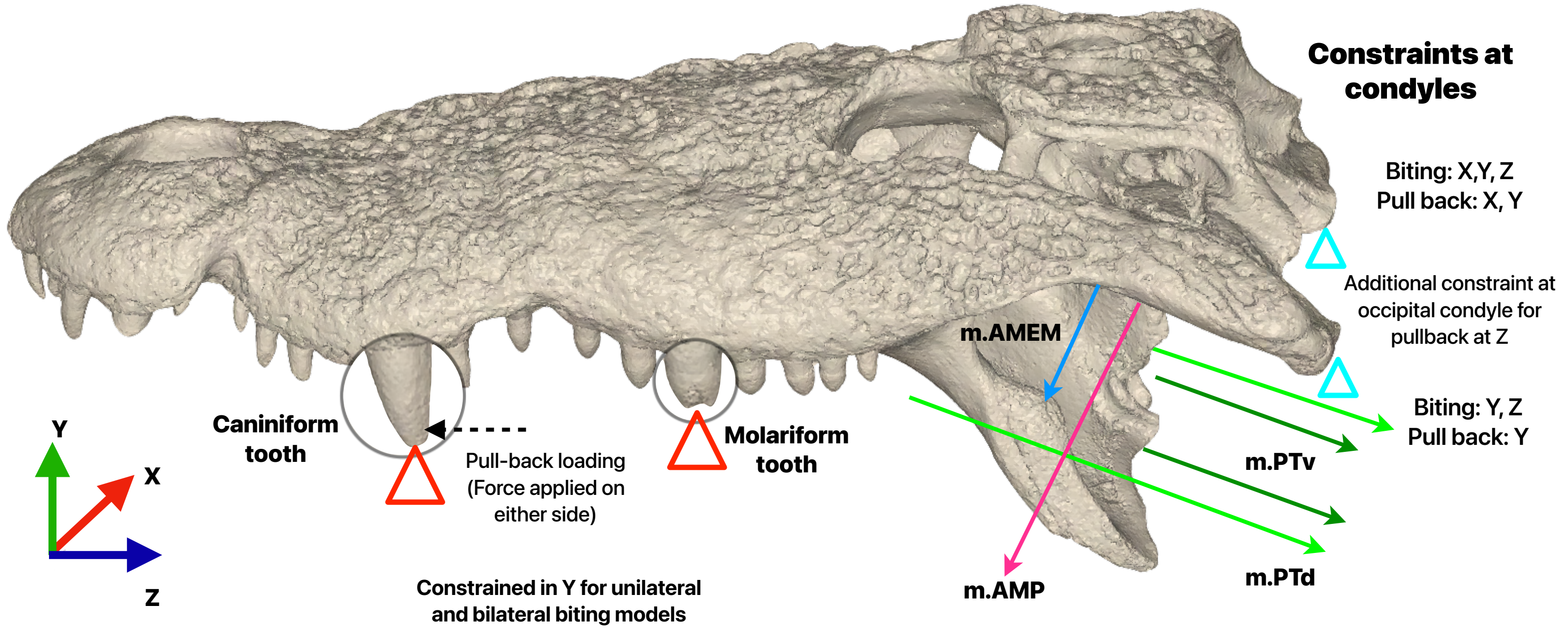


**Figure S3.** Loading conditions applied to the FE models in HyperMesh, shown in *C.niloticus*. Black arrows indicate direction in which teeth are constrained for feeding scenarios. Triangles indicate the boundary conditions applied to the quadrate condyle and teeth. Multi-color arrows represent force vectors, indicating direction of muscle action. Caniniform and Molariform bite points are magnified and highlighted in *C.niloticus*. Global coordinate directions: X = mediolateral, Y = dorsoventral and Z = anteroposterior.

| Taxon | Total Force Applied (in N) | Per Node on each side (in N) |
|---|---|---|
| *Baurusuchus salgadoensis* | 2400 | 1200 |
| *Alligator mississippiensis* | 2000 | 1000 |
| *Montealtosuchus arrudacamposi* | 1000 | 500 |
| *Caipirasuchus paulistanus* | 400 | 200 |
| *Crocodylus niloticus* | 5000 | 2500 |
| *Paleosuchus palpebrosus* | 1000 | 500 |

**Table S3.** Loads applied on the distal side of the caniniform teeth during the pull-back loading.

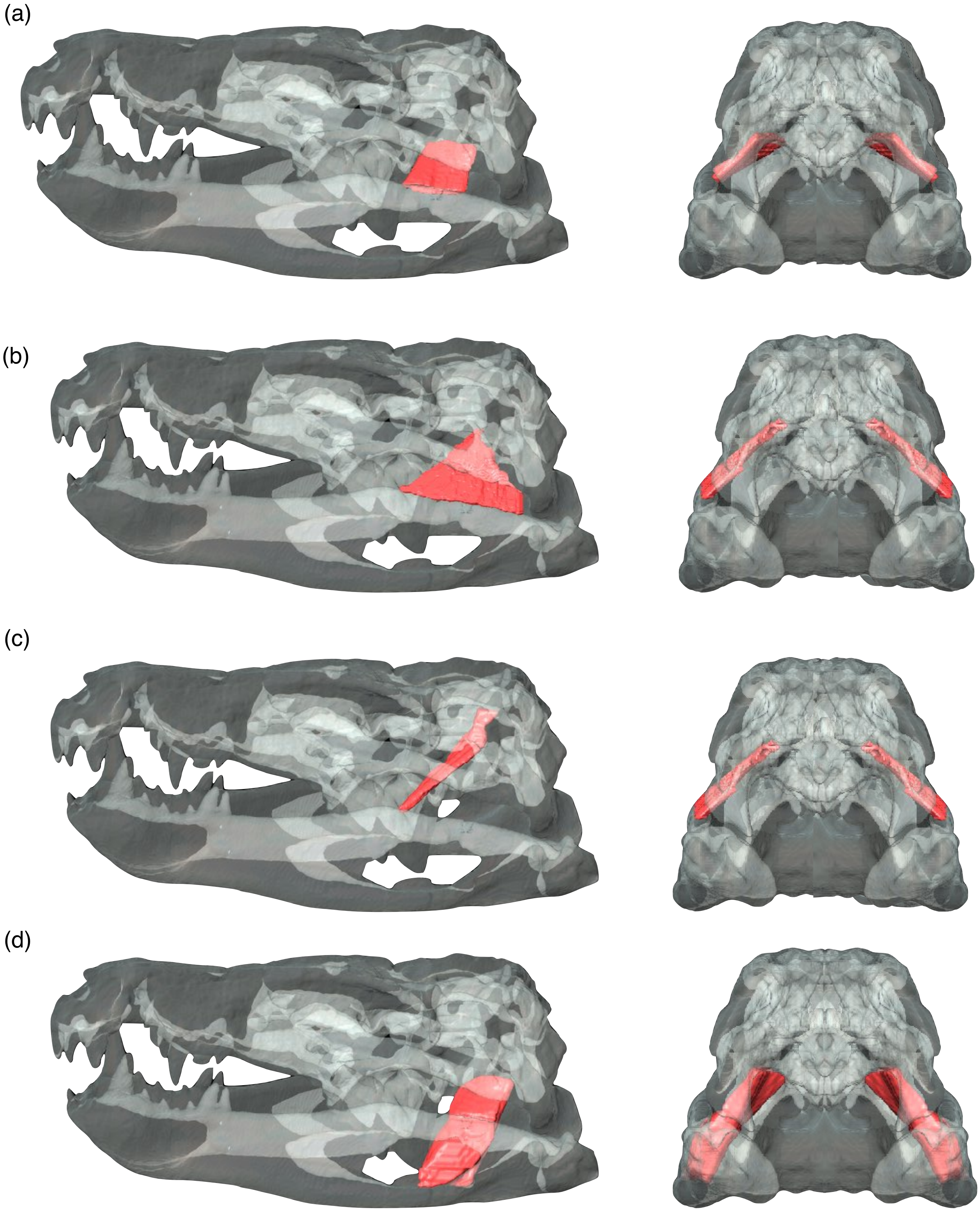


**Figure S4.** Lateral and posterior views of **i**ndividual adductor muscle reconstructions in *Baurusuchus salgadoensis*. (a) m. AMEM, (b) m. AMES, (c) m. AMEP and (d) m. AMP. Bone is rendered transparent.

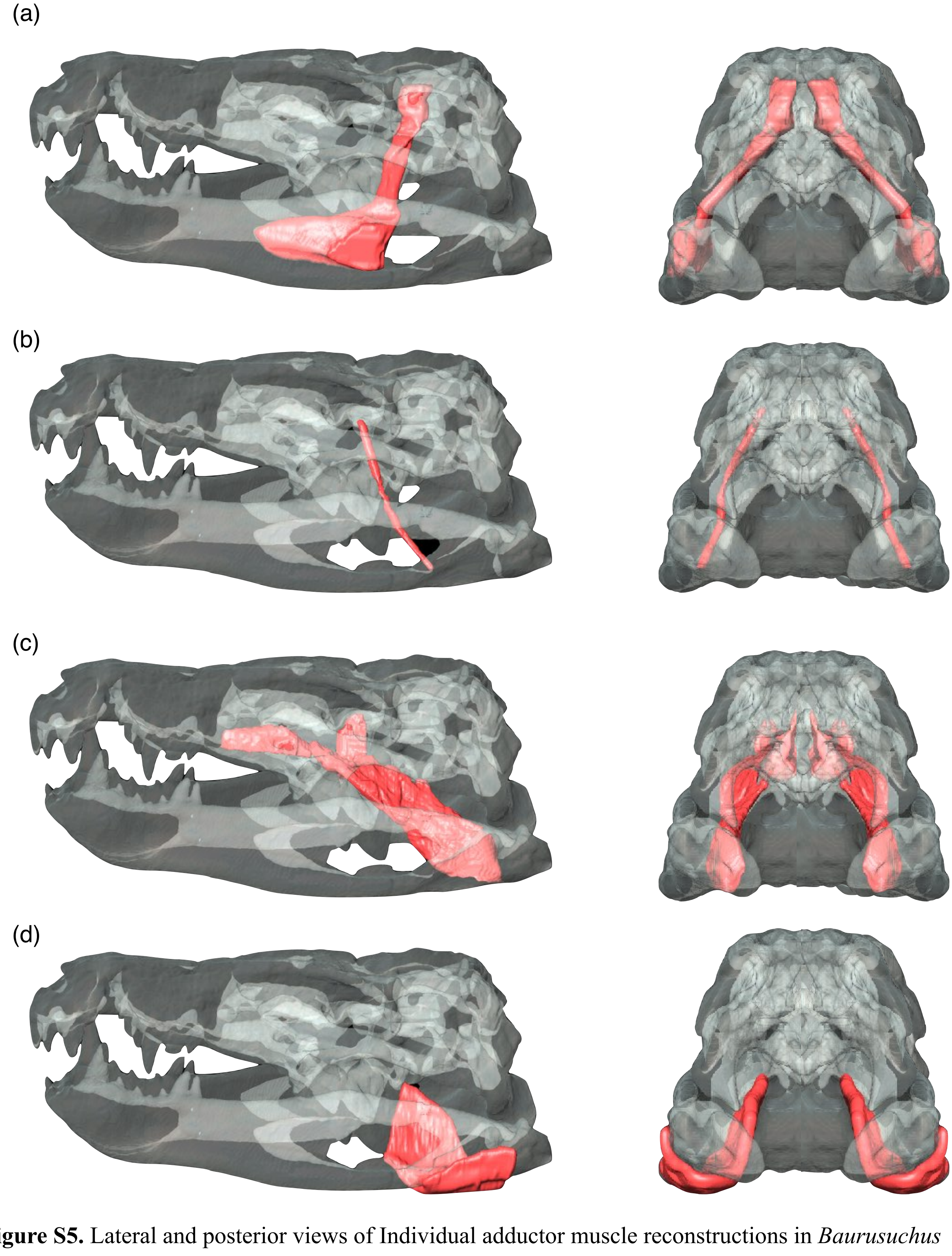


**Figure S5.** Lateral and posterior views of Individual adductor muscle reconstructions in *Baurusuchus salgadoensis*. (a) m. PSTs (including m.IRA and Cartiliago transiliens), (b) m. PSTp, (c) m. PTd and (d) m. PTv. Bone is rendered transparent.

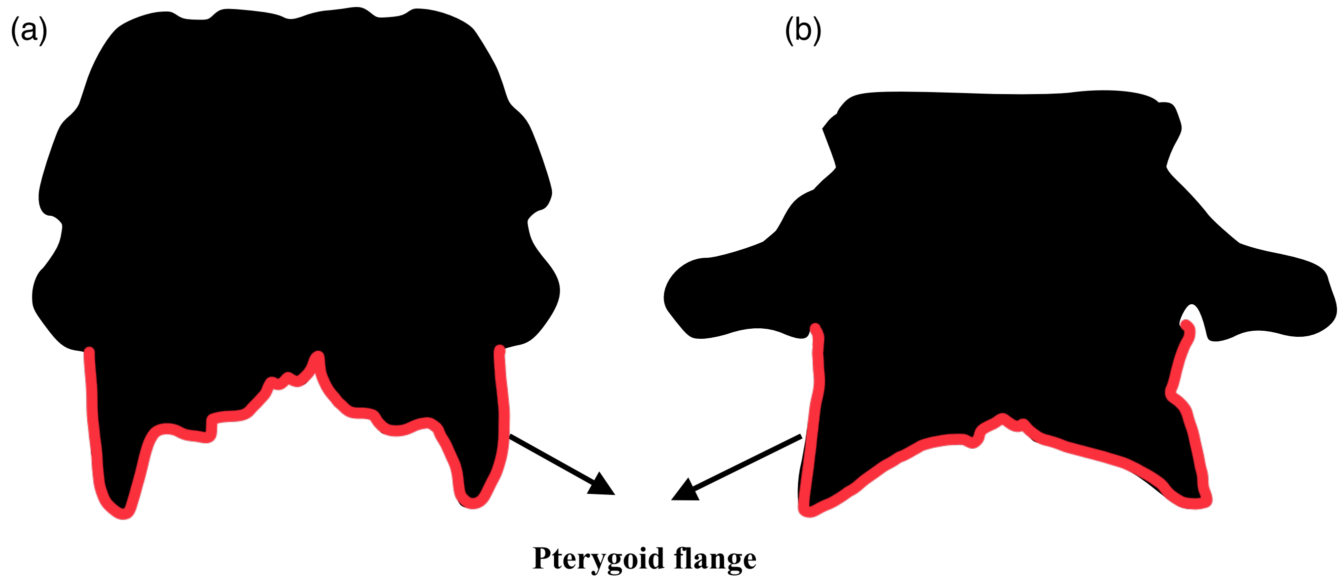


**Figure S6.** Comparison of the pterygoid flanges (highlighted in red), in oreinirostral and platyrostral morphologies. (a) Vertically oriented pterygoids in *Baurusuchus salgadoensis,* and (b) Medially aligned pterygoids in *Paleosuchus palpebrosus*.

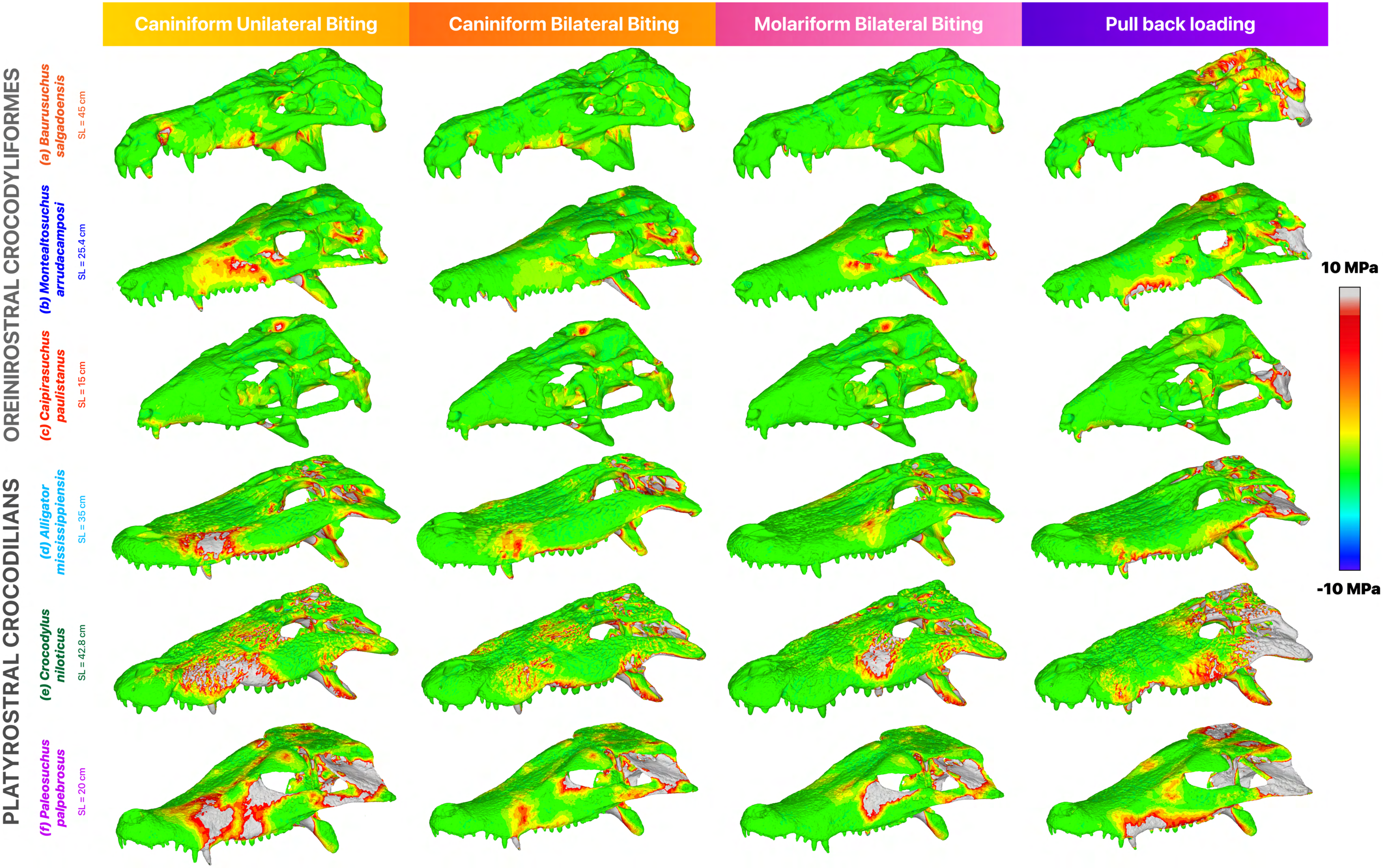
Caniniform Unilateral Biting
Caniniform Bilateral Biting
Molariform Bilateral Biting
Pull back loading
OREINIROSTRAL CROCODYLIFORMES
PLATYROSTRAL CROCODILIANS
(a) Baurusuchus salgadoensis
SL = 45 cm
(b) Montealtosuchus arrudacamposi
SL = 25.4 cm
(c) Caipirasuchus paulistanus
SL = 15 cm
(d) Alligator mississippiensis
SL = 35 cm
(e) Crocodylus niloticus
SL = 42.8 cm
(f) Paleosuchus palpebrosus
SL = 20 cm
10 MPa
-10 MPa

**Figure S7.** From top to bottom: Maximum principal stress distribution in the cranium during various feeding scenarios for (a) *Baurusuchus salgadoensis,* (b) *Montealtosuchus arrudacamposi,* (c) C*aipirasuchus paulistanus,* (d) *Alligator mississippiensis,* (e) *Crocodylus niloticus* and (f) *Paleosuchus palpebrosus.* SL denotes skull length. Warmer colors (yellow to red) indicate regions under higher tensile stress, while cooler colors (green to blue) represent lower or compressive stresses. Areas in grey depict stresses greater than 10 MPa.

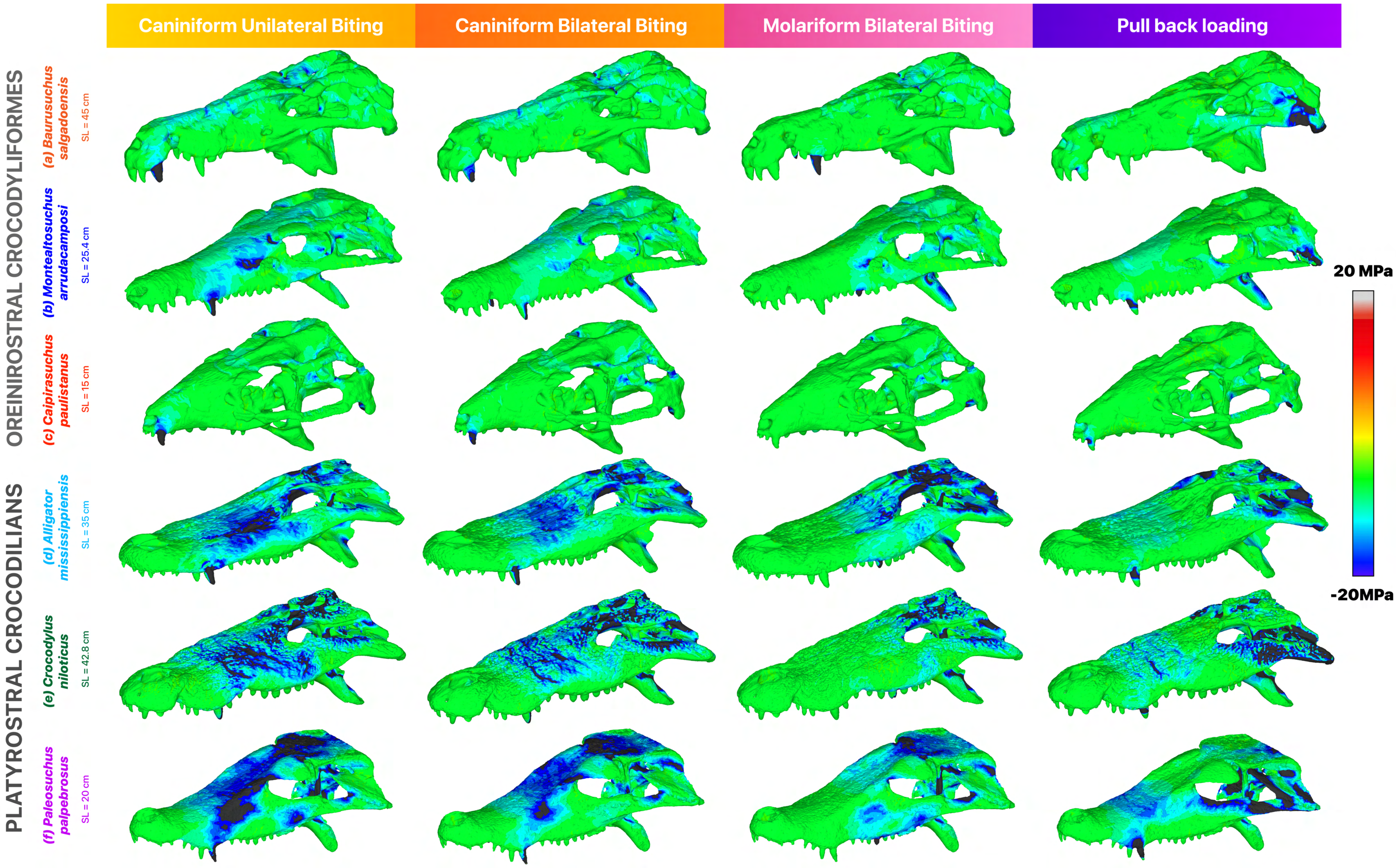

Caniniform Unilateral Biting
Caniniform Bilateral Biting
Molariform Bilateral Biting
Pull back loading
OREINIROSTRAL CROCODYLIFORMES
PLATYROSTRAL CROCODILIANS
(a) Baurusuchus salgadoensis
SL = 45 cm
(b) Montealtosuchus arrudacamposi
SL = 25.4 cm
(c) Caipirasuchus paulistanus
SL = 15 cm
(d) Alligator mississippiensis
SL = 35 cm
(e) Crocodylus niloticus
SL = 42.8 cm
(f) Paleosuchus palpebrosus
SL = 20 cm
20 MPa
-20MPa

**Figure S8.** From top to bottom: Minimum principal stress distribution in the cranium during various feeding scenarios for (a) *Baurusuchus salgadoensis,* (b) *Montealtosuchus arrudacamposi,* (c) C*aipirasuchus paulistanus,* (d) *Alligator mississippiensis,* (e) *Crocodylus niloticus* and (f) *Paleosuchus palpebrosus.* SL denotes skull length. Cooler colors (blue) indicate regions under higher compressive stress, while warmer colors (green to yellow) represent lower compressive or tensile stresses. Areas in grey depict stresses greater than 20 MPa.

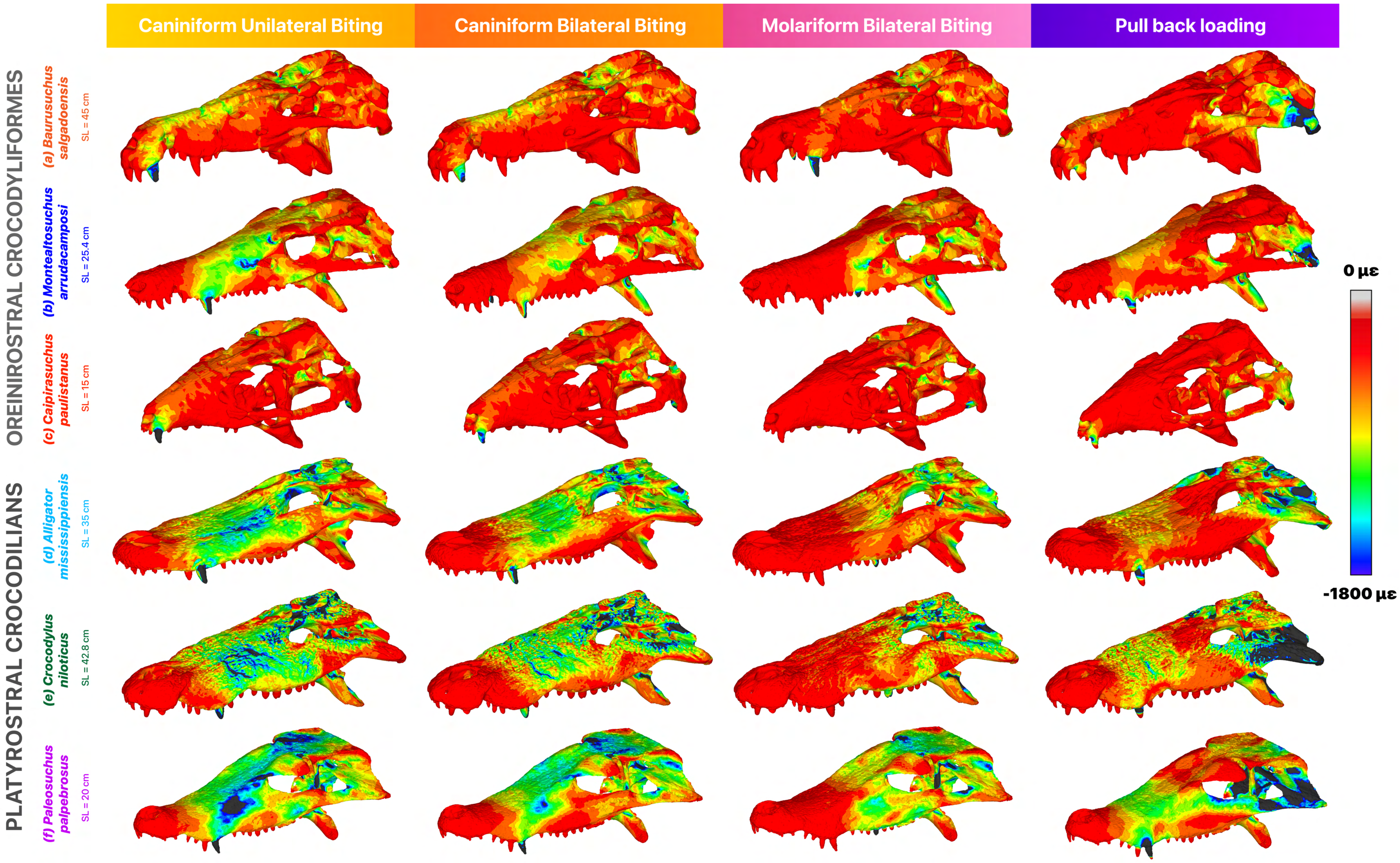
Caniniform Unilateral Biting
Caniniform Bilateral Biting
Molariform Bilateral Biting
Pull back loading
OREINIROSTRAL CROCODYLIFORMES
PLATYROSTRAL CROCODILIANS
(a) Baurusuchus salgadoensis
SL = 45 cm
(b) Montealtosuchus arrudacamposi
SL = 25.4 cm
(c) Caipirasuchus paulistanus
SL = 15 cm
(d) Alligator mississippiensis
SL = 35 cm
(e) Crocodylus niloticus
SL = 42.8 cm
(f) Paleosuchus palpebrosus
SL = 20 cm
0 με
-1800 με

**Figure S9.** From top to bottom: Minimum principal strain distribution in the cranium during various feeding scenarios for (a) *Baurusuchus salgadoensis,* (b) *Montealtosuchus arrudacamposi,* (c) C*aipirasuchus paulistanus,* (d) *Alligator mississippiensis,* (e) *Crocodylus niloticus* and (f) *Paleosuchus palpebrosus.* SL denotes skull length. Cooler colors (blue) indicate regions under higher compressive strain, while warmer colors (green to yellow) represent lower compressive strains approaching zero.

**Table S4.** Image attributions for silhouettes derived from PhyloPic (**www.phylopic.org**), as used in Figure 1.

| **Taxon** | **Artist** | **License and Attribution** |
|---|---|---|
| *Terrestrisuchus gracilis* | Scott Reid | Attribution 3.0 Unported |
| *Hallopus victor* | Jaime A. Headden | Attribution 4.0 International |
| *Protosuchus* | Nobu Tamura, (vectorized by T. Michael Keesey) | Attribution-ShareAlike 3.0 Unported |
| *Epoidesuchus* | Jacob Schick | Attribution 4.0 International |
| *Armadillosuchus arrudai* | Nobu Tamura, vectorized by Zimices | Attribution-ShareAlike 3.0 Unported |
| *Baurusuchus albertoi* | Smokeybjb, vectorized by Zimices | Attribution 3.0 Unported |
| *Sarcosuchus imperator* | Jagged Fang Designs | CC0 1.0 Universal Public Domain |
| *Torvoneustes carpenteri* | Dmitry Bogdanov (vectorized by T. Michael Keesey) | Attribution 3.0 Unported |
| *Alligator mississippiensis* | Ferran Sayol | CC0 1.0 Universal Public Domain |
| *Paleosuchus* | Armin Reindl | Attribution-NonCommercial 3.0 Unported |
| *Gavialis gangeticus* | Jagged Fang Designs | CC0 1.0 Universal Public Domain |
| *Crocodylus porosus* | Smokeybjb | Attribution-ShareAlike 3.0 Unported |

#### R CODE TO CALCULATE 95% MEDIAN STRESS VALUES

```
rm(list=ls())

nile_unilateral <- read.table("/Users/ananthsrinivas/Downloads/Nile_unilateral.txt", sep = "", header = T,na.strings ="", stringsAsFactors= F)

attach(nile_unilateral)

names(nile_unilateral)

n <- 95

bottom <- nile_unilateral[nile_unilateral$SMises < quantile(nile_unilateral$SMises,prob=n/100),] #To remove the top 5% and retain 95% values

mean(nile_unilateral$SMises, na.rm = TRUE) #Mean SMises value

median(nile_unilateral$SMises, na.rm = TRUE) #Median SMises value

median(bottom$SMises, na.rm=TRUE) #Median SMises of top 95% values

bottom1 <- nile_unilateral[nile_unilateral$EMax < quantile(nile_unilateral$EMax,prob=n/100),] #To remove the top 5% and retain 95% values

mean(nile_unilateral$EMax, na.rm = TRUE) #Mean EMax value

median(nile_unilateral$EMax, na.rm = TRUE) #Median EMax value

median(bottom1$EMax, na.rm=TRUE) #Median EMax of top 95% values
```
